# Engagement with the nucleus enhances the efficiency of basement membrane invasion

**DOI:** 10.1101/2025.03.13.643012

**Authors:** Johan d’Humières, Lianzijun Wang, David R. Sherwood, Julie Plastino

## Abstract

Cell invasion through basement membrane (BM) extracellular matrix barriers is important during organ development, immune cell trafficking, and cancer metastasis. Here we study an invasion event, anchor cell (AC) invasion, which occurs during *Caenorhabditis elegans* development. The actin protrusion of the invading AC deforms then disrupts the BM. With live-cell imaging, we observe that the distal end of the actin protrusion contacts the nucleus during wild-type invasion, and deforms the nuclear envelope. Further we show that there is a correlation between invasion efficiency and nuclear contact: under mutant conditions where invasion is reduced, nuclear deformation is diminished. Knocking down the nuclear cytoskeleton protein lamin produces a nuclear positioning defect in the AC and reduces invasion efficiency. In keeping with this, both actin and microtubule-associated nesprins, ANC-1 and UNC-83 respectively, are found on the AC nuclear envelope during invasion, and perturbing their function reduces invasion. CEC-4, a chromodomain protein that attaches chromatin to the nuclear envelope in *C. elegans*, is also found to promote AC invasion. All together these data indicate that the positioning and properties of the AC nucleus are important for BM invasion.

**SUMMARY STATEMENT:** Actin-based membrane protrusions in invading cells apply force to basement membrane (BM) barriers to help break through them. Here we provide evidence that the nucleus also plays a role in optimizing BM invasion.

## INTRODUCTION

Invasion is the process by which cells penetrate through basement membrane (BM) extra-cellular matrix barriers. Invading cells employ active mechanisms, involving actin polymerization and matrix digestion by proteases, in order to create holes in BM as the pore size is too small to allow cells to traverse these dense, cross-linked structures (Castro-Castro et al., 2016; Nürnberg et al., 2011). Indeed discontinuities in BM in patient biopsies are associated with an unfavorable prognosis in many types of cancer (Chang and Chaudhuri, 2019). The role of the nucleus in the initial stages of cell invasion is not clear, although at later stages, the nucleus acts as an impediment during motility in constrained environments, and is actively deformed (Denais and Lammerding, 2014; Patel et al., 2026).

Here we sought to determine the role of the nucleus in BM invasion using a visually accessible *in vivo* invasive event, anchor cell (AC) invasion in *Caenorhabditis elegans*. AC invasion occurs as part of the normal development of the *C. elegans* reproductive system, where the uterine AC generates a large actin protrusion that opens the underlying BM, thus connecting the uterine and vulval epithelial cell layers in order to form the egg-laying apparatus (Sherwood and Sternberg, 2003). AC invasion resembles pathological cancer cell invasion, including the dependence on actin dynamics and proteases for correct invasion (Cáceres et al., 2018; Kelley et al., 2019). Unlike cancer cell invasion, AC breaching of the BM is not followed by migration through the BM gap: the invasive protrusion retracts once invasion is complete, and the AC eventually fuses with its neighbors. In this way, AC invasion allows for the study of how holes are made in BM barriers, without the potentially confounding contributions of subsequent motility.

The mechanical link between the nucleus and the cell cytoskeleton is called Linker of Nucleoskeleton and Cytoskeleton (LINC) complex. In mammals the LINC complex is composed of nesprins and SUN proteins: nesprins are outer nuclear envelope proteins defined by their Klarsicht/ANC-1/Syne homology (KASH) domains, which bind to SUN proteins in the perinuclear luminal space. SUN proteins cross the inner nuclear membrane and bind to lamins in the nuclear interior (Janota et al., 2020). Lamins are nuclear intermediate filaments that form networks under the nuclear envelope, important for the properties and functions of the nucleus (Vahabikashi et al., 2022a). In mammals there are several different isoforms of lamin, which assure different functions (Vahabikashi et al., 2022b). On the cytoplasmic side, different nesprins bind the different components of the cell cytoskeleton (actin, microtubules, intermediate filaments), and thus the LINC complex provides a mechanical link from the cytoplasm to the nuclear interior.

Actin and microtubule cytoskeleton components are well-conserved between mammals and *C. elegans*, but LINC complexes in the worm are somewhat different. First *C. elegans* has a single lamin gene that codes for the protein LMN-1. LMN-1 is generally considered to be a B-type lamin, but since it is the only one, the differences observed in mammals between the roles of A/C-type and B-type are not relevant (Gregory et al., 2023b). Second there are two nesprins expressed in somatic cells in *C. elegans* and one SUN protein: ANC-1 (orthologous to Nesprin-1 and -2), UNC-83 (functionally related to mammalian Nesprin-4 and KASH5) and UNC-84 (orthologous to mammalian Sun1 and Sun2) (Starr, 2019). Both ANC-1 and UNC-83 carry KASH domains, and ANC-1 binds actin filaments, similar to Nesprin-1/2, while UNC-83 interacts with the microtubule cytoskeleton by binding adaptor proteins of the microtubule motors kinesin and dynein (Starr, 2019). Both nesprins interact with the SUN protein UNC-84. The ANC-1/UNC-84 and the UNC-83/UNC-84 LINC complexes are known to be involved in nuclear positioning and anchoring in *C. elegans* (Fridolfsson and Starr, 2010; Malone et al., 1999; Starr, 2019; Starr and Han, 2002; Starr et al., 2001).

In this study we investigate the role of the AC nucleus in BM invasion. Through live cell imaging, mutant analysis and RNAi mediated depletion, we observe that contact between the invasive actin protrusion and the nucleus is important for efficient invasion. We also provide evidence that the LINC complex and heterochromatin anchoring to the AC nuclear periphery promote BM invasion.

## RESULTS AND DISCUSSION

### Nuclear shape and the actin protrusion

AC invasion occurs reproducibly at a specific stage of worm development that can be identified by the stereotyped divisions of the underlying vulval precursor cell (VPC) P6.p (Sherwood and Sternberg, 2003). This allows a large number of invasion events to be observed and quantitative data to be obtained from synchronized worm populations. In wild-type invasion, the AC forms a hole in the BM separating the uterine and vulval epithelial tissues between the P6.p late 2-cell stage (two daughters of P6.p) and the early 4-cell stage (four daughters of P6.p), a period of ∼90 minutes (Figure 1A) (Kenny-Ganzert and Sherwood, 2024). At the early-to-mid 2-cell stage, the BM is intact and actin accumulates at the invasive edge accompanied by BM deformation (Cáceres et al., 2018). By the 4-cell stage, a hole as large as the AC is created in over 98% of worms (N = 62) in the wild-type case, and the actin protrusion fills the opening (Figure 1A).

**Figure 1.**
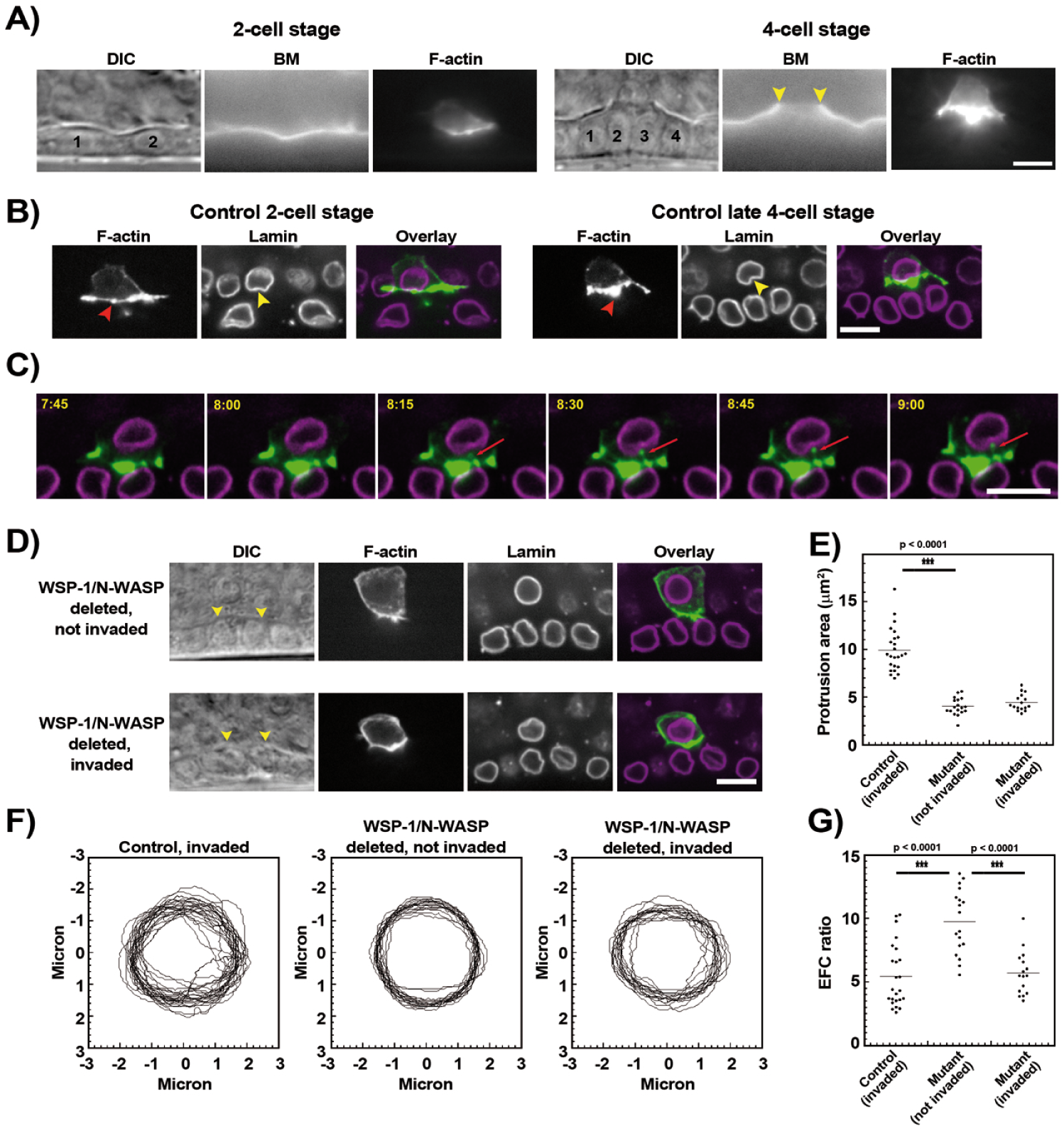
Nuclear shape and the actin protrusion during AC invasion. **A)** AC invasion progression from the P6.p 2-cell, uninvaded stage to the 4-cell stage where invasion is complete, as visualized by the hole in BM fluorescence (yellow arrowheads). Worm strain JUP60. **B)** Invasive actin structures (red arrowheads) deform the AC nucleus (yellow arrowheads) at both the 2-cell and the 4-cell stage in control worms displaying wild-type invasion. Worm strain JUP138. **C)** Still images from a timelapse acquisition. Red arrows indicate an actin structure pushing into the nucleus and deforming it. Time stamps are in minutes:seconds. Worm strain JUP138. **D)** Worms carrying a null mutation in the WSP-1/N-WASP gene *wsp-1*. In most cases **(upper panels)**, the AC does not invade by the 4-cell stage (intact BM indicated by yellow arrowheads on the DIC image), although there are some **(lower panels)** that manage (disrupted BM indicated by yellow arrowheads in the DIC image). In the former case, AC nuclei appear smooth and round, while in the latter case, AC nuclei are deformed like for control animals. Worm strain JUP139. **E)** F-actin protrusion area at the 4-cell stage for control, invaded ACs, and *wsp-1* mutant ACs, both not invaded and invaded. **F)** Overlays of nuclear contours at the 4-cell stage for control, invaded AC nuclei, and *wsp-1* mutant AC nuclei, both not invaded and invaded. All contours are oriented with the invasive front of the AC towards the bottom. **G)** EFC ratios to evaluate nuclear shape irregularity for the contours shown in **F)**. The nuclei of mutant, uninvaded ACs are significantly smoother than those of invaded control ACs and invaded mutant ACs. **A)** epifluorescence microscopy, **B), C)** and **D)** single plane images, spinning disk microscopy of mCherry-labeled laminin (BM), GFP-labeled Lifeact (F-actin) and tagRFP-T-labeled endogenous lamin. See methods for genotypes. The hemispheric plane is shown, i.e. the one with the largest footprint of the nucleus and the protrusion. Scale bars 5 μm. N ≥ 16 for all observations.

During invasion we visualized the AC nucleus using worm strains carrying fluorescently-labeled, endogenous nuclear lamin. Although reduced brood size and embryo viability have been linked to tagging endogenous lamin with full-length GFP (Bone et al., 2016; Gregory et al., 2023b; Link et al., 2018), AC invasion was completely wild-type (96% invasion, N = 47). We observed that, at earlier stages where actin assembled in the form of foci that depressed the underlying BM (Hagedorn et al., 2013), foci corresponded to dents in the AC nuclear lamin contour (Figure 1B). Even more striking, the large actin protrusion of later stages was clearly in contact with the nucleus as evinced by substantial depressions in the nuclear envelope (Figure 1B). Live imaging of actin dynamics and nuclear lamin during AC invasion showed that the formation of nuclear dents was indeed correlated with the growth of actin structures abutting the nucleus (Figure 1C and Supplementary Movie S1).

### Nuclear deformation and invasion

To gain insight into the functional significance of protrusion-nucleus contact, we examined a worm strain that was null for the gene of the single homolog of the N-WASP protein, WSP-1 (Withee et al., 2004). N-WASP is an actin polymerization activator that catalyzes the activity of the Arp2/3 complex, one of the main actin polymerization nucleation factors responsible for actin structure assembly in moving and protruding cells (Blanchoin et al., 2014). Previous work had shown that the invasive actin protrusion in the AC is much reduced in *wsp-1* null animals, and successful invasion occurs in only about 20% of 4-cell stage worms (Cáceres et al., 2018; Lohmer et al., 2016). We evaluated nuclear contours in ACs of *wsp-1* null animals that had not invaded at the 4-cell stage, and we observed that the nuclei were smoother and rounder than in the control conditions (Figure 1D, upper panels). As previously observed (Cáceres et al., 2018), the size of the actin protrusion was halved in mutant ACs that had not invaded (Figure 1E). We overlaid uninvaded and control nuclear contours for a whole population of worms, and worms lacking WSP-1 that did not display invasion at the 4-cell stage had smoother AC nuclei that superimposed more neatly than control worms, which had erratic nuclear outlines (Figure 1F). We quantified this effect using the elliptical Fourier analysis approach, as previously used to analyze irregular nuclear shapes ((Tamashunas et al., 2020) and references therein). Briefly this method fits the nuclear contour with a series of diminishing ellipses, and then outputs the EFC ratio, which characterizes how well the nuclear shape was fit by the first ellipse. As such the EFC ratio is larger for more regular shapes. Indeed, the mutant, uninvaded ACs had nuclei with an average EFC ratio that was approximately double the control AC nuclei (Figure 1G).

This result linked a smooth nuclear shape to inefficient invasion. To test this further, we took advantage of biological variability and the fact that some *wsp-1* null worms succeeded in invading. These worms had nuclei that were deformed (Figure 1D, lower panels) despite the fact that the AC actin protrusions were small in size, similar to uninvaded ACs (Figure 1E).

In fact, overlay analysis and EFC ratio quantification revealed that mutant ACs that succeeded in invading had nuclei as deformed as control animals (Figure 1F and G). These data were consistent with the idea that invasion happened in those *wsp-1* animals where the little actin that was present was organized in such a way as to contact the nucleus, as evidenced by the observed deformations. Overall from this part, we concluded that the actin protrusion contacted and deformed the nucleus during efficient invasion.

### Role of the nucleocytoskeleon in AC invasion

Given this correlation, we next wondered whether the nucleocytoskeleton was important for AC invasion. We targeted by RNAi the most upstream component of the LINC complex, the lamin protein LMN-1. 100% of treated AC nuclei displayed a patchy, non-continuous lamin shell, and total fluorescent intensity on the nuclear envelope was reduced by 70% as compared to control nuclei, evaluated by performing *lmn-1* RNAi in fluorescent lamin strains (Supplementary Figure S1). Nevertheless RNAi against *lmn-1* in a control strain gave no significant effect on invasion efficiency, even when assessed at the early 4-cell stage, where invasion was less robust (Figure 2A). This was not particularly surprising as it was known that LMN-1 was a stable protein on one hand (J. Liu et al., 2000), and that a little went a long way: a homozygous *lmn-1* deletion obtained from a heterozygous, balanced strain showed no morphological or movement defects, presumably due to the maternal load of LMN-1 although this was present at levels undetectable by Western blot (Haithcock et al., 2005).

**Figure 2.**
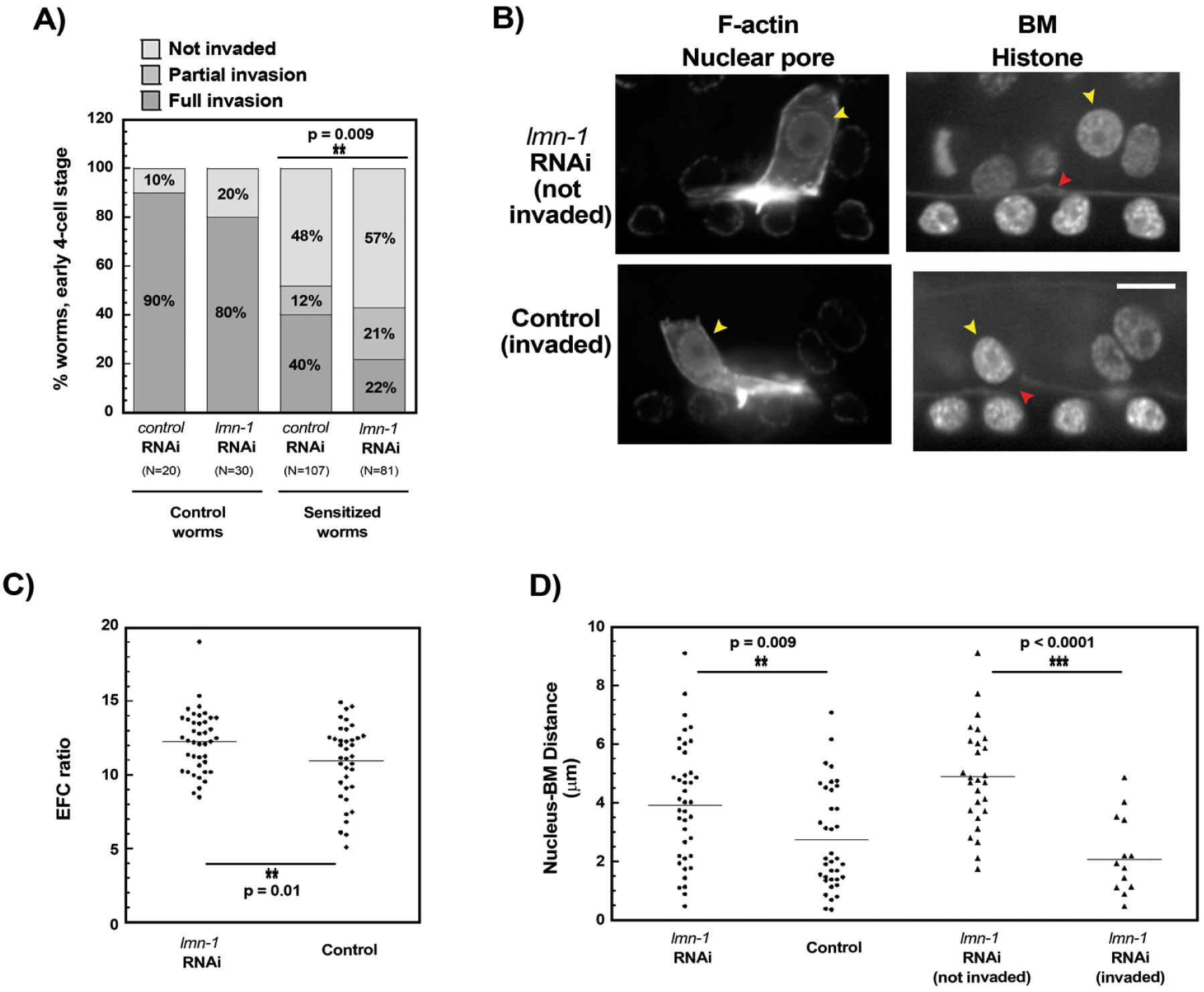
Role of nuclear lamin in AC invasion. **A)** Quantification of invasion phenotypes under *lmn-1* RNAi: full invasion, partial invasion (when a hole in BM is present, but smaller than normal), not invaded (when the BM is intact). Invasion is assessed at the early 4-cell stage. Control worms are strain JUP60. Sensitized worms are protease-deficient, strain JUP65. See methods for genotypes. Control RNAi is with empty vector *L4440*. **B)**, **C)** and **D)** Nuclear contours and positioning under *lmn-1* RNAi as compared to control RNAi in sensitized worms. **B)** Nuclear pores and F-actin are visible in the same channel, but AC nuclei can be distinguished (yellow arrowheads), and their shape quantified by tracing nuclear contours defined by nuclear pore signal. Distances are measured between the edge of the AC nucleus, as seen with the histone label (yellow arrowheads) and the BM, either intact or its former position in the case of invasion (red arrowheads). Single hemispheric-plane images. Spinning disk microscopy of GFP-labeled Lifeact (F-actin), GFP-labeled NPP-21 (nuclear pore), mCherry-labeled laminin (BM) and mCherry-HIS-58 (histone). Worm strain JUP150. See methods for genotypes. Scale bar 5 μm. **C)** EFC ratios for *lmn-1* RNAi-treated nuclei versus control RNAi treatment measured by tracing nuclear contours defined by nuclear pores. N ≥ 37. **D) Circles:** Nucleus-BM distances measured under *lmn-1* RNAi treatment and control treatment. N ≥ 37. **Triangles:** The *lmn-1* treatment data is then further divided into not invaded and invaded populations showing that long distances are strongly correlated with unsuccessful invasion. N ≥ 14 in each pool.

With these caveats in mind, we turned to a sensitized worm strain that had been used successfully in the past to reveal cryptic mechanisms, a strain lacking the two main matrix metalloproteases (MMPs) involved in AC invasion (Kelley et al., 2019). Indeed AC invasion is a robust process, and compensating mechanisms can mask the functional consequences of applied perturbations. For example RNAi against *arx-2*, a subunit of the Arp2/3 complex, the main actin polymerization nucleating factor driving AC invasion, has only a moderate effect on its own, but when performed in a MMP-deficient background, the removal of *arx-2* drastically reduces AC invasion, especially at early stages (Kelley et al., 2019). Similarly in a protease-deficient background, *lmn-1* RNAi produced a significant invasion defect at the early 4-cell stage, with only 22% successful invasion, versus 40% for the control treatment (Figure 2A).

To examine the nuclear envelope during invasion under lamin RNAi conditions, we used a strain carrying fluorescent nuclear pores (Thomas et al., 2023). We observed that *lmn-1* RNAi-treated nuclei had slightly larger EFC ratios than control nuclei (Figure 2B and 2C), indicating that RNAi-treated nuclei were less deformed, although EFC ratios were large in all conditions as nuclei were very smooth at this early stage of invasion where actin structures were small. More notably, *lmn-1* RNAi treatment produced a significant defect in the positioning of the AC nucleus, with larger distances observed under *lmn-1* disruption (Figure 2B and 2D, filled circles). Indeed when the *lmn-1* RNAi data was separated into invaded and not invaded pools, there was a very significantly larger average distance between the nucleus and the BM in the non-invaded case as compared to the invaded case (Figure 2D, filled triangles). We concluded that attachment of the nucleus to the cytoskeleton and presence of the nucleus in proximity to the BM was important for invasion efficiency.

### Role of the cytoskeleton-nucleus link in AC invasion

Since *lmn-1* RNAi gave such dramatic positioning defects, we next looked at the nesprin proteins ANC-1 and UNC-83, as these proteins connect the nucleus to the actin and microtubule cytoskeletons, respectively. Indeed since the SUN protein, UNC-84, requires lamin for correct nuclear envelope localization, unlike mammalian SUN proteins (Gregory et al., 2023a; Hasan et al., 2006; Lee et al., 2002), our *lmn-1* RNAi would be expected to perturb the connection of both ANC-1 and UNC-83 to the nucleus.

Using strains with the endogenous loci of *anc-1* and *unc-83* fluorescently labeled via CRISPR, we first confirmed that these nesprins were present in the AC at the time of invasion. ANC-1 was found on the nuclear envelope in AC, but appeared also in the cytoplasm, indicating association with the endoplasmic reticulum, as per previous reports in other *C. elegans* cell types (Figure 3A) (Hao et al., 2021; Starr and Han, 2002). UNC-83, on the other hand, was found exclusively associated with the nuclear envelope in a punctate pattern, also as observed previously (Figure 3B) (Starr et al., 2001).

**Figure 3.**
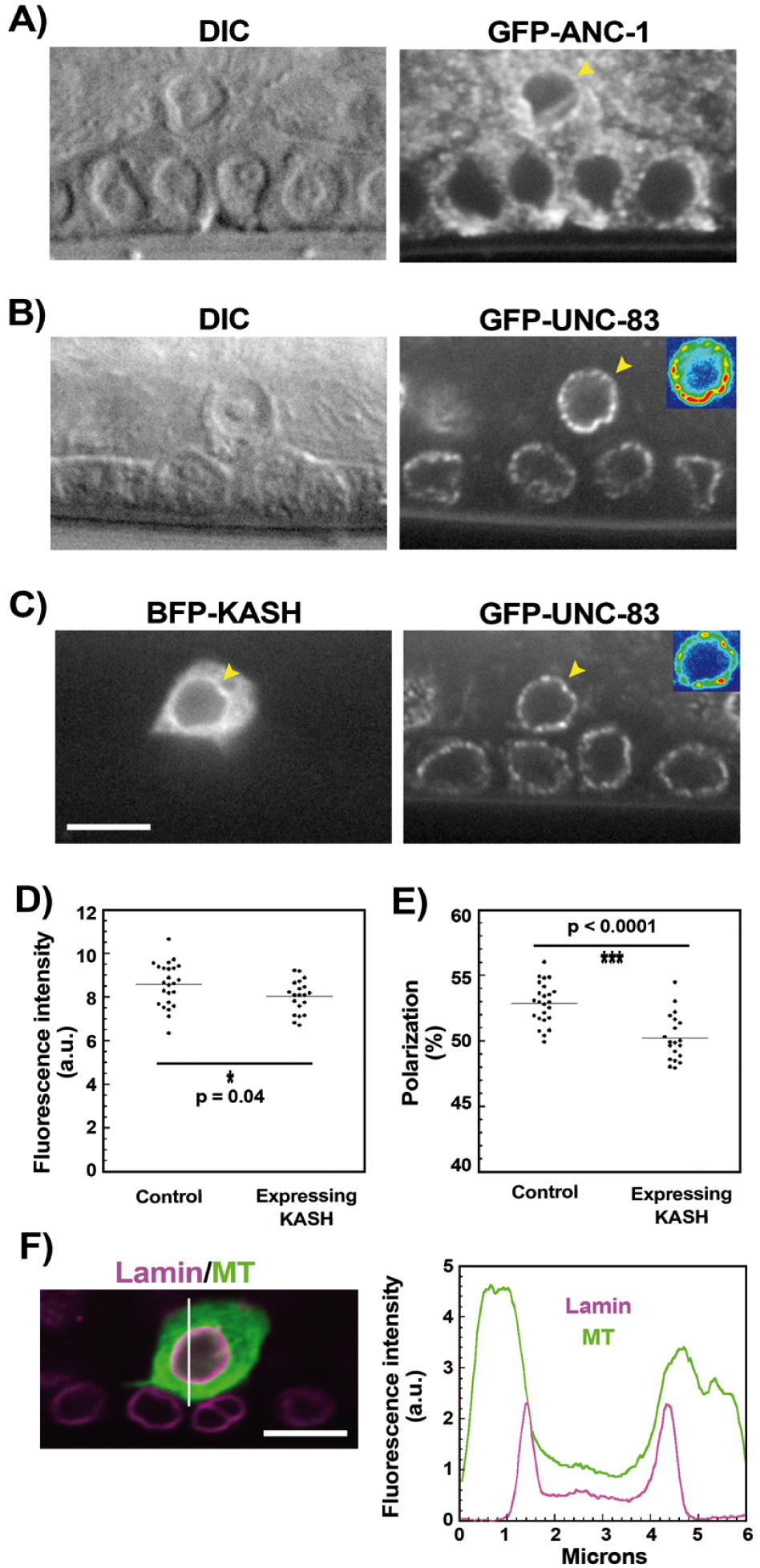
Probing the molecular attachment between the AC nucleus and the cytoskeleton. **A)** Fluorescent labeling of endogenous full-length ANC-1 on the N-terminus by CRISPR shows that ANC-1 is expressed in the AC at the time of invasion (4-cell stage), and concentrated on the nuclear envelope (yellow arrowhead) although also present in the cytoplasm. Worm strain MCP341. **B)** Fluorescent labeling of endogenous full-length UNC-83, with GFP inserted near the KASH domain by CRISPR, shows this nesprin in a punctate pattern on the AC nucleus (yellow arrowhead) and slightly brighter near the invasive front as emphasized by the inset heatmap. Worm strain UD473. **C)** The dominant negative KASH construct, expressed in the AC, is enriched at the nuclear envelope (yellow arrowhead, BFP channel), but only slightly reduces UNC-83 signal at the nuclear envelope (yellow arrowhead, GFP-UNC-83 channel), although polarization is lost as highlighted by the inset heatmap. Worm strain JUP149. **D)** Quantification of reduction of UNC-83 signal (fluorescence intensity) on the AC nuclear envelope upon KASH expression. **E)** Quantification of loss of polarized UNC-83 signal on the AC nuclear envelope upon KASH expression. Data is represented as %, i.e., the fluorescent signal found in the half-contour facing the BM divided by the total fluorescence of the entire contour. **F)** Visualization of fluorescently-labeled endogenous lamin on the nucleus (magenta) and microtubules (green), observed via expression of EMTB-GFP in the AC. The white line indicates the position of the linescan shown on the right, where the peak of MT fluorescence basal with respect to the lamin peak is larger than that observed at the back of the nucleus. Worm strain NK3622. See methods for all genotypes. DIC coupled with single plane spinning disk images. In all cases the hemispheric plane is shown, i.e. the one with the largest footprint of the AC nucleus. All scale bars 5 μm.

Since both nesprins were present on the AC nuclear envelope, we tried to disrupt both at once by a dominant negative approach that has been extensively used in mammalian cells: overexpression of a KASH domain that saturates the SUN binding sites on the nuclear envelope, displacing nesprin proteins and thereby rupturing the nucleus-cytoskeleton link (Lombardi et al., 2011; Stewart-Hutchison et al., 2008). In the AC we overexpressed the ANC-1 KASH domain as it binds more strongly than the UNC-83 KASH domain to the SUN domain of UNC-84 (Jahed et al., 2019). Animals bearing KASH in their ACs did not display aberrant invasion as compared to control animals (Table 1). However very little UNC-83 appeared to be displaced by KASH expression, despite the fact that KASH was correctly enriched on the nuclear envelope (although also abundant in the cytoplasm) (Figure 3C and 3D). Intriguingly KASH perturbation called attention to the fact that the UNC-83 signal was slightly enriched on the hemisphere of the nucleus directed toward the invasion site, and this polarization was lost upon KASH expression (Figure 3B, 3C, inset heatmaps and Figure 3E). We could not assess if ANC-1 was polarized like UNC-83, and/or displaced by KASH expression as the nuclear envelope and cytoplasmic signals of GFP-ANC-1 were impossible to distinguish. UNC-83 polarization was reminiscent of the nesprin polarization observed in mammalian cell nuclei navigating through microfluidic constrictions, where actin was shown to pull nesprins to the front of the nucleus and concentrate them there (Davidson et al., 2020). UNC-83 is a nesprin that associates with microtubule motors (Fridolfsson et al., 2010; Meyerzon et al., 2009), and indeed we observed that microtubules were enriched between the AC nuclear envelope and the invasive edge, visualized by AC-specific expression of fluorescently-labeled EMTB, a domain of ensconsin that binds along the sides of microtubules (Figure 3F) (Kenny-Ganzert et al., 2026; Richardson et al., 2014).

**Table 1:** Effect of KASH expression on AC invasion at the 4-cell stage.

|  | RNAi treatment | % full invasion | N |
| --- | --- | --- | --- |
| Control<sup>1</sup> | None | 98%<sup>a</sup> | 62 |
| KASH-expressing<sup>2</sup> | None | 97% | 58 |
| *anc-1* null<sup>3</sup> | *unc-83* RNAi | 89%<sup>a</sup> | 54 |

<sup>1</sup>JUP60, <sup>2</sup>JUP124, <sup>3</sup>JUP116. See methods for genotypes.
<sup>a</sup> Significantly different: p = 0.05

From this result, it appeared that KASH overexpression was only effective at displacing a labile microtubule-attached subpopulation of UNC-83 from the nuclear envelope. To disrupt nesprin function more thoroughly, we performed *unc-83* RNAi in an *anc-1* null mutant, and obtained a small but significant reduction in AC invasion (Table 1). Combining the *lmn-1* RNAi findings, nesprin localization and polarization observations and this RNAi result, we concluded that molecular attachments between the AC cytoskeleton and nesprins on the AC nucleus contributed modestly to efficient invasion.

### Role of heterochromatin tethering in AC invasion

Since the AC nucleus appeared to be submitting to mechanical stresses, we wondered if heterochromatin could play a role as it is known to protect nuclei during cell movements and shape changes in other systems (Kalukula et al., 2022; Stephans et al., 2018). A recent paper on nuclear migration though constricted spaces in *C. elegans* shows a role for tethering of H3K9-methylated chromatin to the nuclear envelope via CEC-4, a chromodomain protein (Gregory et al., 2025). Of note altering *cec-4* releases H3K9-methylated heterochromatin from the nuclear membrane without affecting global transcription (Gonzalez-Sandoval et al., 2015). Furthermore *lmn-1* RNAi has no effect on CEC-4, and CEC-4 and LMN-1 signals do not colocalize on the nuclear envelope (Gonzalez-Sandoval et al., 2015), so *cec-4* perturbation gave us a parallel way of testing the role of nuclear properties in AC invasion.

We therefore knocked down *cec-4* by RNAi, first in a strain carrying fluorescently-labeled endogenous CEC-4 in order to evaluate RNAi efficiency (Gonzalez-Sandoval et al., 2015). The effect was all or nothing in the AC: approximately half the worms had no detectable CEC-4-mCherry under *cec-4* RNAi (N = 28) as compared to one out of 22 for the control RNAi (Supplementary Figure S2). Given this incomplete RNAi, we evaluated *cec-4* RNAi directly in the sensitized strain at the early 4-cell stage, as we had done for *lmn-1* RNAi. There was no effect on invasion (Figure 4A). This was not particularly surprising as Gregory et al. had found that the effect of perturbing *cec-4* on P-cell nuclear migration was only revealed when the LINC complex was also disrupted, perhaps because both CEC-4 and LINC complex contribute to protecting the nucleus from mechanical insults (Gregory et al., 2025). We therefore repeated *cec-4* RNAi in a version of the sensitized strain that additionally expressed the dominant negative KASH domain, and observed a substantial reduction in invasion (Figure 4A). Evaluation of nuclear contours revealed a small but significant increase in EFC ratio under *cec-4* RNAi in this background, indicating that nuclei were less deformed (Figure 4B).

**Figure 4.**
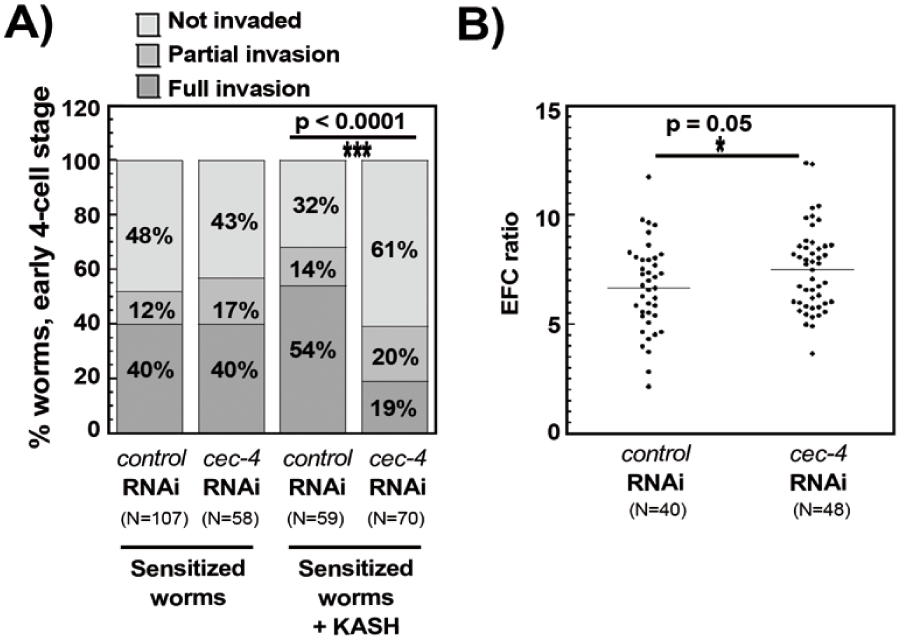
Role of chromatin attachment to the nuclear envelope in AC invasion. **A)** Quantification of invasion phenotypes under *cec-4* RNAi: full invasion, partial invasion (when a hole in BM is present, but smaller than normal), not invaded (when the BM is intact). Invasion is assessed at the early 4-cell stage. Sensitized worms are protease-deficient, strain JUP65. Sensitized worms expressing KASH are strain NK3475. Control RNAi is with empty vector *L4440*. **B)** Nuclear deformations are quantified in protease-deficient worms expressing KASH and also fluorescently-labeled endogenous lamin, strain NK3439. See methods for genotypes.

## Conclusion

Overall our results indicate, in multiple contexts, that ineffective invasion is associated with a smooth nucleus, whether the actin cytoskeleton, nucleus attachment to the cytoskeleton or heterchromatin tethering to the nuclear envelope are being perturbed. In particular the phenotypic diversity of the N-WASP deletion allows us to compare AC invasion in identical (reduced) actin conditions, and observe that invasion is associated with deformed AC nuclei. This suggests that a suboptimal amount of actin is sufficient to assure invasion when organized properly to abut and deform the nucleus. Unfortunately we would need super-resolution microscopy to see filament organization, something we have tried in the AC previously with little success due to the extreme denseness of the invasive protrusion (Dreier et al., 2019). Our live imaging is consistent with the idea that growing actin structures deform the nucleus with their distal end during invasion. This is reminiscent of nuclear indentations observed in invading human cancer cells in culture (Revach et al., 2015).

Interfering with the LINC complex connection reduces AC invasion efficiency, and altering heterochromatin tethering exacerbates this effect, as also observed in another cell shape change event during *C. elegans* development (Gregory et al., 2025). It is possible that this is due to synergy between independent effects: untethering heterochromatin that fragilizes the AC nucleus on one hand and weakening the cytoskeleton connection and perturbing nuclear positioning in the AC on the other hand.

One scenario that is consistent with our observations is that the invasive actin protrusion in the AC pushes down on the BM, braced on the distal side by the nucleus, which is held in place by nesprin interaction with the actin and microtubule cytoskeletons. In this context it is important to note that polymerization of actin networks in cells can produce 1 to 10 kPa of pressure, depending on cell type (Abraham et al., 1999; Brunner et al., 2006), and that we have previously reported that the AC exerts about 30 nN on the BM (Cáceres et al., 2018), which translates to kPa range of pressures depending on the protrusion size. Nuclei have an average elastic modulus in about the same range (Fisher et al., 2020; Krause et al., 2013; H. Liu et al., 2014). Together this suggests that the actin within the invasive protrusion deforms the nucleus.

All together the data point toward a role for the nucleus in AC invasion, perhaps as a brace for the invasive actin protrusion, and associate a smooth nucleus with reduced invasive efficiency. It remains to be seen if this correlation is confirmed in other systems where cells are making holes in BM, and if nuclear deformation coinciding with BM disruptions could be a useful parameter for assessing metastatic potential in patient biopsies.

## SUPPLEMENTARY FIGURE AND MOVIE LEGENDS

**Supplementary Figure S1.**
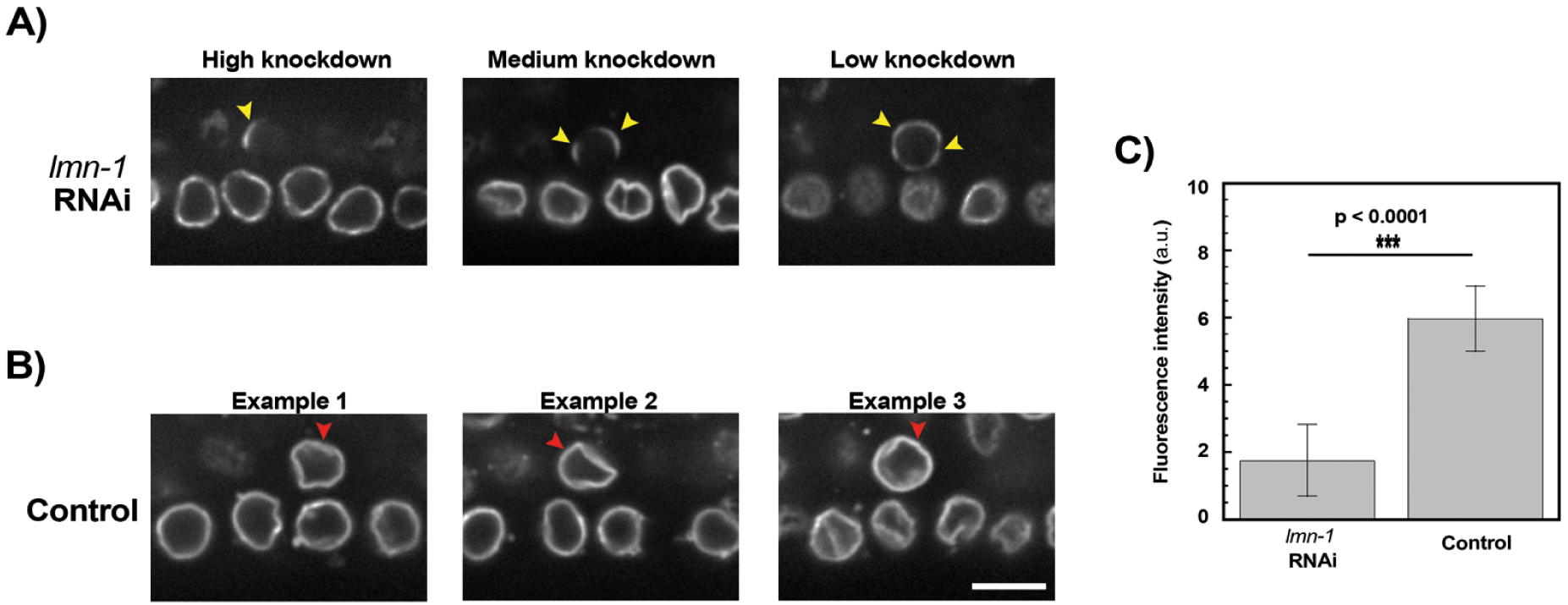
Assessment of *lmn-1* RNAi efficiency. **A)** Examples of high, medium and low knockdown efficiency observed with *lmn-1* RNAi. Residual lamin patches on AC nuclei are indicated with yellow arrowheads. VPC nuclei are visible beneath the AC as RNAi is observed to be very ineffective in the VPCs in most cases. **B)** Examples of control RNAi with the *L4440* empty vector. Intact lamin shells on AC nuclei are indicated with red arrowheads. **C)** Quantification of fluorescence intensity. The integrated fluorescence at the nuclear envelope in the hemispheric plane of each nucleus was measured and averages ± standard deviations are shown. N ≥ 9. Spinning disk microscopy. Worm strain JUP138. See methods for genotype. Scale bar 5 μm.

**Supplementary Figure S2.**
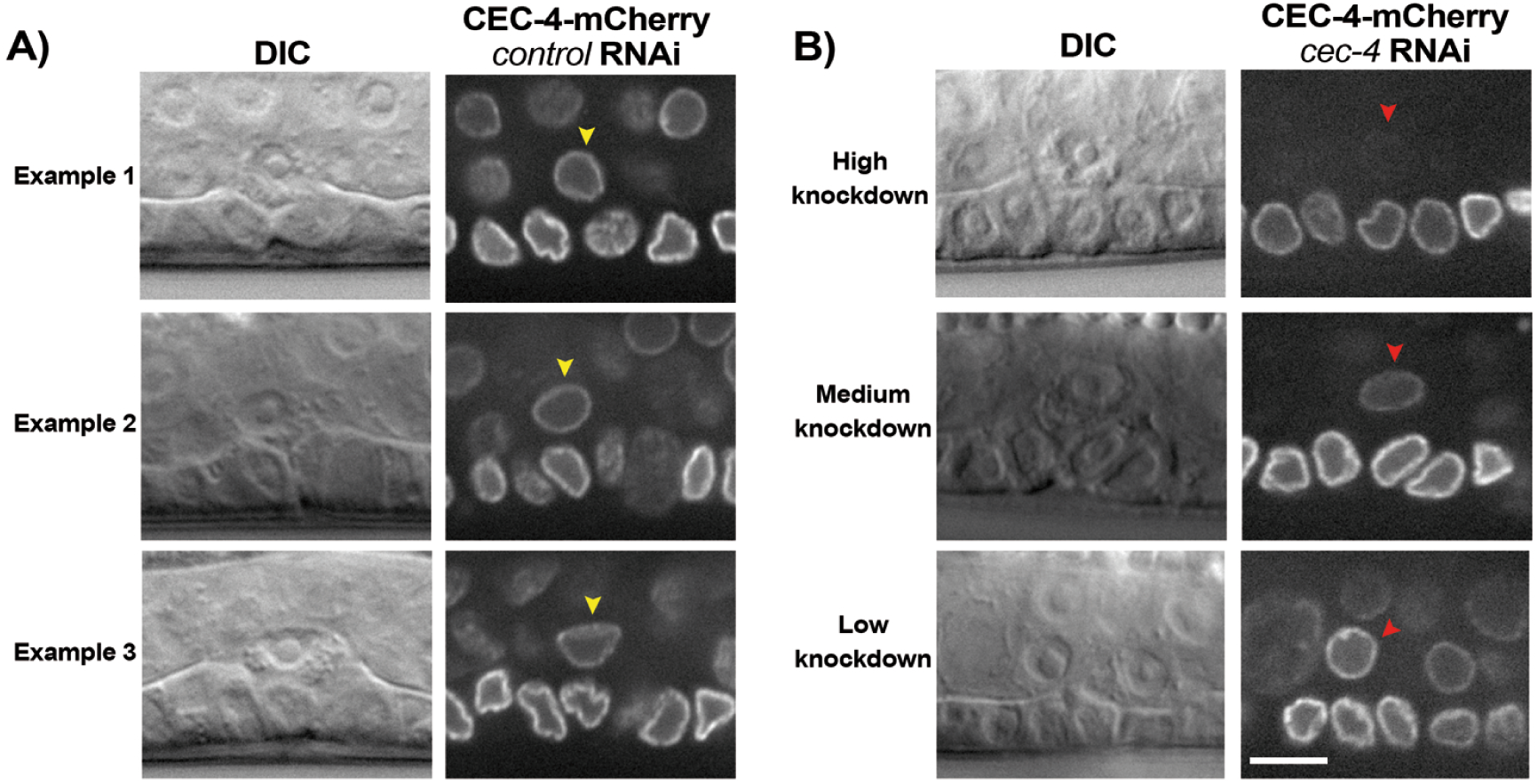
Assessment of *cec-4* RNAi efficiency. **A)** Examples of control RNAi with the *L4440* empty vector. The AC nuclei display a ring of CEC-4 at the nuclear periphery (yellow arrowheads). **B)** Examples of the different CEC-4 signals observed in the AC (red arrowheads) upon *cec-4* RNAi. Knockdown efficiency is highly variable with only about half of the sample displaying high knockdown in the AC (top panels). Single plane spinning disk images. In all cases the hemispheric plane is shown, i.e. the one with the largest footprint of the AC nucleus. Worm strain GW849. See methods for genotype. All scale bars 5 μm.

**Supplementary Movie S1 The distal side of the actin protrusion contacts the AC nucleus and deforms it.** Movie to accompany still images in Figure 1C. Actin structures are seen to form, abut the AC nucleus and deform it. Autoscaling for the intense brightness of the invasive front makes it appear that there is an empty space between some parts of the actin protrusion and the AC nucleus, but movements in the actin are mirrored by deformations of the nucleus, indicating that they are in contact. Time stamps are in minutes:seconds. GFP-labeled Lifeact (green) and tagRFP-T-labeled endogenous lamin (magenta). Worm strain JUP138. See methods for genotype. Single plane spinning disk of the hemispheric plane, i.e. the one with the largest footprint of the AC nucleus and the actin protrusion. Scale bar 5 μm.

**SUPPLEMENTARY TABLE 1:**
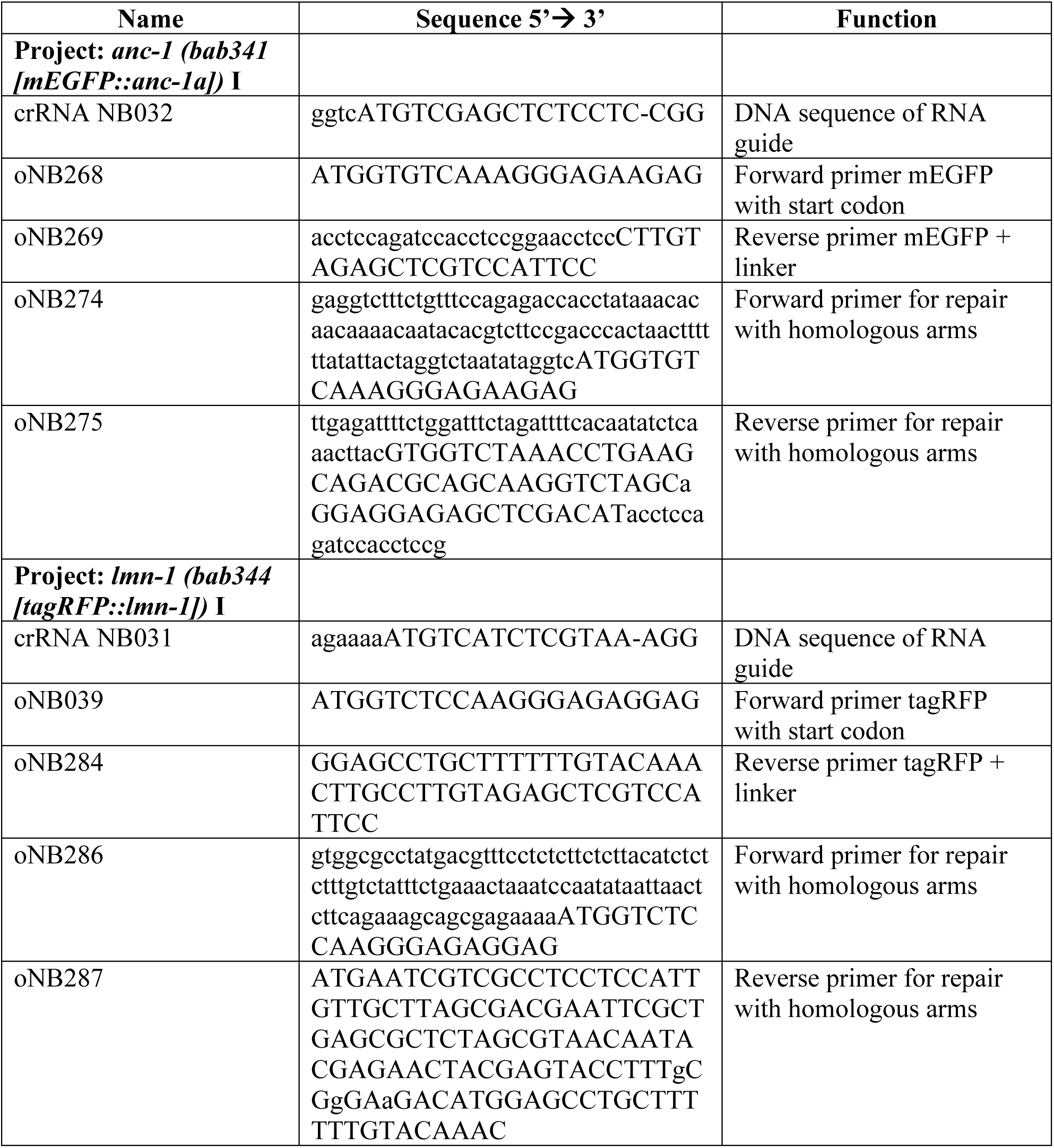
Oligo sequences used to generate the CRISPR strains.

## MATERIALS AND METHODS

### Constructs, alleles and worm strains

The alleles *qyIs127 [Plam-1::lam-1::mCherry]*, *qyIs242 [Pcdh-3::Lifeact::GFP]*, *zmp-1(cg115)* III and *zmp-6 (tm3073) III* are described in previous studies (Cáceres et al., 2018; Kelley et al., 2019). Allele *qyIs636 [lin-29p::EMTB::GFP::unc-54 3’UTR + unc-119(+)]* is described in (Kenny-Ganzert et al., 2026). The strains CB3440 *anc-1(e1873)* I and NG324 *wsp-1(gm324)* IV were provided by the *Caenorhabditis* Genetics Center. CRISPR/CAS9-genome edited strain UV117 *lmn-1(jf98[lmn-1::GFP])* I was a kind gift from Verena Jantsch (Link et al., 2018). CRISPR/CAS9-genome edited strain BN1062 *npp-21 (bq1[npp-21::GFP]); bqSi189[pBN13(unc-119(+) lmn-1p::mCherry::his-58)]* II with fluorescent nuclear pores and histone was from Peter Askjaer (Thomas et al., 2023). UD473 *unc-83 (yc26[unc-83::GFP::kash+LoxP])* V, a form of *unc-83* with an internal GFP label introduced by CRISPR/CAS9, was gifted by Daniel Starr (Bone et al., 2016; Jahed et al., 2019). The MosSCI insertion strain GW849 *gwSi17 [cec-4p::cec-4::WmCherry::cec-4 3’UTR]* II was provided by Jan Padeken and Susan Gasser (Gonzalez-Sandoval et al., 2015).

CRISPR/CAS9-genome edited strain MCP341 *anc-1 (bab341 [mEGFP::anc-1a])* I and strain MCP344 *lmn-1 (bab344 [tagRFP::lmn-1])* I were generated at the SEGiCel facility (Université Lyon 1, UMS3421, Lyon, France) following the method reported in (Dokshin et al., 2018). For *anc-1a*, the longest, full-length form was labeled on its N-terminus with a monomeric form of enhanced GFP containing a syntron, followed by GASGASGAS linker before the start ATG. For *lmn-1*, tagRFP-T with syntrons followed by the linker ASLYKKAGS was introduced on the N-terminus of the gene coding for LMN-1, as had been done for the GFP version (Link et al., 2018). Oligonucleotide sequences used to create these strains are detailed in Supplementary Table 1. Of note an N-terminally-labeled form of endogenous *unc-83* was also created by CRISPR/CAS9, but it was poorly expressed and not correctly localized to the nuclear envelope, so UD473 was used instead.

The gene for the dominant negative KASH construct (allele *sybIs5165*) was synthesized by Eurofins and consisted of the tag-BFP gene with syntrons followed by a GGSGGGSGG linker and then the neck region, transmembrane domain and the luminal domain of ANC-1 as defined in (Hao et al., 2021). This construct was recombined into Gateway cloning vector pDONR221. The promoter sequence *Pzmp-1* and the enhancer element *pes-10* were amplified from *pBS-zmp-1p-pes-10-spGFP1-10* as done in (Hagedorn et al., 2009), and introduced into Gateway cloning vector pDONR[P4-P1R]. The entry vectors were recombined along with the *unc-54 3′UTR* (gift of G. Seydoux; Addgene plasmid #17253: pCM5.37) into the destination vector pCFJ150 - pDESTttTi5605[R4-R3] (gift from Erik Jorgensen Addgene plasmid # 19329). The construct was injected into DP38 *unc-119(ed3)* and integrated by Suny Biotech. The strain was outcrossed at least twice before use.

Crossing different alleles gave the following worm strains in addition to the ones already described:

JUP60 *qyIs127 [Plam-1::lam-1::mCherry]; qyIs242 [Pcdh-3::Lifeact::GFP]*

JUP65 *qyIs127 [Plam-1::lam-1::mCherry]; qyIs242 [Pcdh-3::Lifeact::GFP] ; zmp-1(cg115) III; zmp-6 (tm3073) III*

JUP116 *anc-1(e1873) I; qyIs127 [Plam-1::lam-1::mCherry]*

JUP124 *sybIs5165 [Pzmp-1-pes-10::BFP::KASH::unc-54 3’UTR + unc-119(+)]; qyIs127 [Plam-1::lam-1::mCherry]; qyIs242 [Pcdh-3::Lifeact::GFP]*

JUP138 *qyIs242 [Pcdh-3::Lifeact::GFP]*; *lmn-1 (bab344 [tagRFP::lmn-1])* I

JUP139 *wsp-1(gm324)* IV; *qyIs242 [Pcdh-3::Lifeact::GFP]*; *lmn-1 (bab344 [tagRFP::lmn-1])* I

JUP149 *sybIs5165 [Pzmp-1-pes-10::BFP::KASH::unc-54 3’UTR + unc-119(+)]; unc-83 (yc26[unc-83::GFP::kash+LoxP])* V

JUP150 *npp-21 (bq1[npp-21::GFP]); bqSi189[pBN13(unc-119(+) Plmn-1::mCherry::his-58)]* II; *qyIs127 [Plam-1::lam-1::mCherry]; qyIs242 [Pcdh-3::Lifeact::GFP] ; zmp-1(cg115) III; zmp-6 (tm3073) III*

NK3439 *qyIs242 [Pcdh-3::Lifeact::GFP]*; *lmn-1 (bab344 [tagRFP::lmn-1])* I; *sybIs5165 [Pzmp-1-pes-10::BFP::KASH::unc-54 3’UTR + unc-119(+)]; zmp-1(cg115) III; zmp-6 (tm3073) III*

NK3475 *sybIs5165 [Pzmp-1-pes-10::BFP::KASH::unc-54 3’UTR + unc-119(+)]; qyIs127 [Plam-1::lam-1::mCherry]; qyIs242 [Pcdh-3::Lifeact::GFP] ; zmp-1(cg115) III; zmp-6 (tm3073) III*

NK3622 *qyIs636 [Plin-29::EMTB::GFP::unc-54 3’UTR + unc-119(+)]; lmn-1 (bab344 [tagRFP::lmn-1])* I

### Worm cultivation and RNAi treatment

Strains were grown at room temperature on agar plates containing NGM growth media and seeded with *Escherichia coli* OP50 strain. Worms were synchronized by bleach and starved to arrest at the L1 stage, and then fed OP50 at room temperature until AC invasion was scored. RNAi was performed by feeding bacteria producing target dsRNAs to larvae at room temperature. RNAi plates contained 2 mM IPTG and the HT115 cultures containing the different RNAi probes were brought to 2 mM IPTG before spotting on plates. Phenotypes were scored at either the 4-cell stage or the early 4-cell stage, as specified. Empty *L4440* was used as a negative control. The probe for *cec-4* was obtained from the Vidal ORF RNAi library (Rual et al., 2004). The probes for *lmn-1* and *unc-83* were obtained from the Ahringer RNAi library distributed by Source Bioscience (Kamath and Ahringer, 2003).

### Imaging

Worms were anesthetized in 30-50 μL of 10 mM (0.2%) levamisole for 10-15 min, and then mounted on 4.5% noble agar pads. Samples were sealed with melted VALAP (vaseline, lanolin, paraffin 1:1:1 w/w/w). Invasion was evaluated on an inverted epifluorescence Zeiss microscope equipped with DIC optics, using a 100x objective. Confocal image acquisition was performed at room temperature with an upright confocal spinning disk microscope from Roper/Zeiss using a 100x 1.46 OIL DIC ALPHA PL APO (UV) VIS-IR objective and a CoolSNAP HQ2 camera controlled by Metamorph software. DIC optics were used and laser lines at 405 nm, 491 nm and 561 nm. Time lapse imaging of the JUP138 strain expressing fluorescently tagged LMN-1 and LifeAct was conducted on a Zeiss Axio Imager microscope equipped with a Yokogawa CSU- W1 spinning-disk confocal system and controlled using Micro-Manager software v2.0.1. Images were acquired with a Zeiss 100× Plan-Apochromat 1.4 NA oil-immersion objective and a Hamamatsu ORCA-Quest qCMOS camera. Fluorophores were excited using 488-nm and 561-nm lasers. Slide preparation followed a previously described method (Kelley et al., 2017). A single z slice was captured at each time point at the AC central plane and images were collected at 15-second intervals.

### EFC ratio analysis

Masks for nuclear contours were made by thresholding. These masks were fed into a custom Matlab script, a kind gift of Alice Williart (Matthieu Piel laboratory, Institut Curie) based on the original script developed in (Tamashunas et al., 2020). The number of harmonics, N, was taken as 25.

### Image analysis

In order to quantify *lmn-1* knock-down, tag-RFP-T total intensity on the AC nuclear envelope was quantified by thresholding on the hemispheric plane of the AC nucleus. For the knock-down, the intensity of nuclear patches was summed. For the control, the intensity at the nuclear envelope was quantified by subtracting the inner fluorescence from the total. For obtaining nuclear contours under *lmn-1* RNAi, the NPP-21-GFP (nuclear pore) signal was used. The dotted nature made it impossible to trace automatically, so the dots were connected by hand to give nuclear outlines. For nucleus-BM distances, the red channel was observed where both the nucleus (histone) and the BM were visible. Lines were drawn from the edge of the nucleus to the BM or where the BM would be if not invaded. Lines followed the shape of the AC to capture the mispositioning in both the dorsal-ventral and in the anterior-posterior directions. For quantification of UNC-83 signal with and without KASH, contours were drawn by hand (punctate fluorescence made thresholding unreliable) with a 10-pixel wide line, and average intensities were logged at each position along the contour. These values were summed to give the total fluorescent intensity of the contour. To calculate polarization, the total fluorescent intensity of the front half of the AC nucleus (toward the BM/VPCs) was divided by the total fluorescent intensity of the entire contour.

### Statistical analysis

For comparison of % phenotypes with RNAi, a chi-squared test was employed to generate p values. All other p values calculated by Student t-test. p ≤ 0.05 is taken as significant.

## ACKNOWLEDGEMENTS

Spinning disk imaging was performed at the Cell and Tissue Imaging Platform (member of France–BioImaging, ANR-10-INBS-04) of the Genetics and Developmental Biology Department (UMR3215/U934) of Institut Curie. Some worm strains were obtained from the *Caenorhabditis* Genetics Center, which is funded by NIH Office of Research Infrastructure Programs (P40 OD010440). Some CRISPR strains made in this study were generated by SEGiCel (SFR Santé Lyon Est CNRS UAR 3453, Lyon, France) with the support of CNRS and IBiSA. D.R.S. acknowledges financial support by the National Institutes of Health (Grant R35GM118049). JdH acknowledges the Ministère de l’Enseignement Supérieur et de la Recherche for PhD funding. J.P. acknowledges financial support from the Fondation ARC (Grant PJA 20191209604) and the Human Frontiers Science Program Organization (Grant RGP0026/2020).

